# The Euler Characteristic Transform Enables Classification of Complex Plant Shapes and Prediction of Leaf Venation from Blade Geometry

**DOI:** 10.64898/2026.04.13.718293

**Authors:** Yemeen Ayub, Sarah McGuire-Scullen, Sarah Percival, William N Weaver, Nischal Karki, Wahiba Yahiaoui, Karina Astudillo-Pavón, Alejandra Barrios, Jill C Check, Joel Colchado-López, Benedikt A. Dolgikh, Deborah V. Espinosa-Martínez, Qiuyi Fu, Kevin Manuel Galvan-Lara, J. Noé García-Chávez, Santiago Garcia-Rios, Chloe N Grabb, Geisson E. Guadir-Lara, John C. Hawkins, Chandler L. Hendrickson, Asia T Hightower, Jimena J. Hurtado-Olvera, Sepideh Kianian, James Lennon, Zhuohao Li, Joy Li, Brent Lieb, Jinxin Lin, Patricio López-Sánchez, Mariana Luna-Alvarez, Coral Martínez-Martínez, Aura Montemayor-Lara, Nicholas A Moreno, Idowu Arinola Obisesan, Oscar Pérez-Flores, Carlos Pimentel-Ruiz, Eduardo Pineda-Hernández, Connor Purvis, Luke Sharpe, Safa Smail, Farnaz Tajik, Joshua A Temple, Mitchell A Ticoras, Jacob K. Trusky, Judson J. Van Wyk, Diana Villarreal-Huerta, Elena Xinyi Wang, Nathan Willey, Julia Zheng, Beronda L. Montgomery, Annabel Romero-Hernandez, Emily B Josephs, Zoë Migicovsky, Luis Diaz-Garcia, Aman Y Husbands, Stephen A Smith, Ziane Laiadi, Robert VanBuren, Alejandra Rougon-Cardoso, Elizabeth Munch, Daniel H Chitwood

## Abstract

**Rationale:** Quantifying and predicting plant morphology is central to understanding development and evolution, yet many plant forms lack homologous features required for traditional morphometrics. We apply the Euler Characteristic Transform (ECT), an injective descriptor from topological data analysis, to encode 2D plant shapes. The ECT converts contours into image-like representations that preserve shape information while enabling deep learning.

**Methods:** We computed ECTs for large datasets of leaf and pavement cell shapes and used convolutional neural networks (CNNs) for classification. We also trained CNNs to approximate the inverse mapping, predicting leaf shape masks from radial ECTs.

**Key results:** ECT-based models achieved high classification accuracy, surpassing previous approaches on millions of herbarium-derived leaves. Notably, grapevine leaf venation was predicted from blade geometry alone, demonstrating that vascular structure is encoded in the outline.

**Main conclusion:** The ECT provides a compact, information-preserving representation of biological shape that integrates naturally with deep learning. It enables both accurate classification and predictive reconstruction, revealing latent morphological information and offering new opportunities to study plant form across scales.

## Introduction

Geometric morphometrics is a powerful approach for quantifying and visualizing shapes (Bookstein 1992). Applied analysis of shapes in biology has primarily focused on animals or objects that have determinate forms. In a biological context (distinct from its formal mathematical meaning), the concept of homology, or features that share a descent from a common ancestor, both underlies and constrains modern morphometric methods. Homologous features are used to generate corresponding landmarks, and by extension anchor pseudo-landmarks not based on homology (Bookstein 1997; Hightower et al., 2026), that allow for Procrustean-based methods that minimize a distance metric between samples (Gower 1975; Kendall 1984). Homology underlies outline-based methods like Elliptical Fourier Descriptors (Kuhl and Giardina 1982; Iwata et al. 1998) as well, as prominent shared features between samples impact the resulting harmonic series. Homology between samples permitting these methods enables the visualization of theoretical shapes using inverse functions and the ability to analyze and pinpoint relevant features.

While biological homology is always present, it is more cryptic in some lineages than others, and its inaccessibility limits the application of geometric morphometric approaches as intended. A case in point is plants: sometimes leaves have abundant corresponding features (Viscosi and Cardini 2011), as in the case of palmate leaves, and in grapevine or maracuyá (*Passiflora* spp.), for example, we have applied geometric morphometric approaches successfully (Chitwood and Otoni 2017; Chitwood et al. 2025). In broader comparisons across all lateral organs, biological homology is limited to the base and tip of leaves at best (Hightower et al. 2026). Corresponding features likely exist, but because of the iterative and indeterminate nature of plant development, they are embedded in serially homologous structures (Friedman and Diggle 2011), such as leaflets within compound leaves, which require advanced modeling approaches that are difficult to generalize across broad taxonomic groups (Balant et al. 2024). Other plant shapes entirely lack identifiable homologous features or even orientation outside their original context, including the convoluted features of pavement cells (Vőfély et al. 2019) or root or shoot architecture (Prusinkiewicz 1986).

In cases where geometric morphometrics cannot be applied, we have turned to topological data analysis (TDA) approaches. TDA methods use the mathematical field of topology to measure the shape of data (Munch 2017; Amézquita et al. 2020). A filter function (also known as a lens function) is a real valued function in the input shape. Tools from TDA study topological features in the data across different thresholded subsets of the shape based on this function. In TDA, persistent homology encodes topological features, such as connected components, loops, or voids. For example, one can compute persistence on roots using a geodesic distance function from the root base to measure branching patterns in root (Li et al. 2017) and inflorescence architecture (Li et al. 2019). Another tool from TDA, the mapper graph, uses distance based clustering on level sets of a filter function to visualize underlying data structures as a graph (where a graph consists of vertices and edges), which we have applied to gene expression across the flowering plants (Palande et al. 2023) and to leaf shape changes across the heteroblastic series (Percival et al. 2024). The focus of this paper, the Euler Characteristic Transform (ECT), has been proven to completely encode the information of a shape, meaning no two distinct graphs can have the same ECT and thus the transform preserves all shape information rather than summarizing or discarding it (Turner et al. 2014). In practice, the ECT works by sampling a finite number of directions and computing the Euler characteristic 𝜒 (which in the graph case is defined as the number of vertices minus the number of edges) of sublevel sets of the height function in each direction. The ECT therefore tracks how connected components appear and merge as the shape is scanned at each threshold; by concatenating the information from each direction the ECT can be viewed as an image, in which the vertical axis represents thresholds, the horizontal axis represents sampled angles, and the pixel values encode the Euler characteristic (Munch 2025). One way to think about the ECT is that the shape is scanned from multiple angles as if illuminated from different directions, producing a representation that encodes how structure accumulates across both direction and scale. When data has orientation, the corresponding directional axes between samples allow direct comparison between the features of ECTs, allowing specific features contributing to differences between groups to be identified, as we previously showed with three dimensional X-ray computed tomography data of barley seeds (Amézquita et al. 2022).

Here, we use the ECT to create machine learning inputs for classification and prediction of morphological features across plant species. On large datasets of leaf and pavement cell shapes, we demonstrate its effectiveness as an input to convolutional neural networks (CNNs) for classification tasks. The ECT enables high classification accuracy for data with diverse shape features and no corresponding (biological) homology that has been impossible to meaningfully interpret using previous geometric approaches that rely on these properties. The ECT also achieves a higher classification rate across plant families than previously observed (Li et al., 2018). Using a semantic segmentation CNN, we show that the original shape mask can be predicted from its ECT. Finally, using synthetically augmented data for training, the shape mask and ECT of the blade of grapevine leaves reasonably predicts the veins, demonstrating that the information required to reconstruct vasculature is embedded within the leaf outline. Overall, we introduce a computational mathematical technique to the analysis of plant morphological data at different scales that enables analysis of previously intractable morphological problems and integrates seamlessly with modern machine learning approaches.

## Materials and Methods

### Euler Characteristic Transform (ECT)

The ECT summarizes a shape by “scanning” it from many directions and, at each step of the scan, measuring the topology of the shape by calculating the Euler characteristic. For a chosen direction, we sweep a line in that direction and, at each position, compute the difference between the number of vertices and number of edges present in the portion of the object that lies below the line. Specifically, for a 2D shape, stacking these counts across all thresholds (rows) and all directions (columns) produces a compact, image-like representation of the shape.

For intuition, we show the ECT with example visualizations (**Figures 1-2**). We begin with 2D closed contour outlines extracted from paper cutouts taken from *The Parakeet and the Mermaid* (1952) by Henri Matisse, used here for educational purposes (**Figure 1**). These outlines exemplify characteristics common in plant shape data that traditional morphometrics cannot accommodate: 1) variable numbers of non-homologous, repetitive structures within a class, 2) deeply lobed or branching forms, and 3) lack of consistent orientation (Li et al. 2018). Each outline is first converted into a graph with an indeterminate number of vertices and edges (**Figure 1A**). The corresponding ECT image is a pixel grid where the vertical axis represents thresholds, the horizontal axis represents angles, and each pixel encodes the Euler characteristic. We calculate the ECT in this manuscript using the recently released “ect” Python module (Ayub et al. 2026). This encoding uniquely allows rotation in the original shape to appear as horizontal translation in the ECT image (McGuire 2024; Munch 2025; **Figure 1B**). By adding redundant padding on each side (regions “A” and “B” in **Figure 1C**), the ECT image can wrap into a cylinder (**Figure 1D**). Combined with typical zero-padding at the top and bottom, this cylindrical structure gives ECT images orientation-invariant properties when analyzed with CNNs. Given the nonlinear relationships between ECT images, we applied manifold learning techniques to visualize sample similarities. For purposes of alignment, we first applied principal component analysis (PCA; Pearson 1901), followed by nonlinear dimensionality reduction using t-distributed stochastic neighbor embedding (t-SNE; Maaten and Hinton 2008; **Figure 1E**), which effectively separates these complex shapes into respective groups.

**Figure 1:**
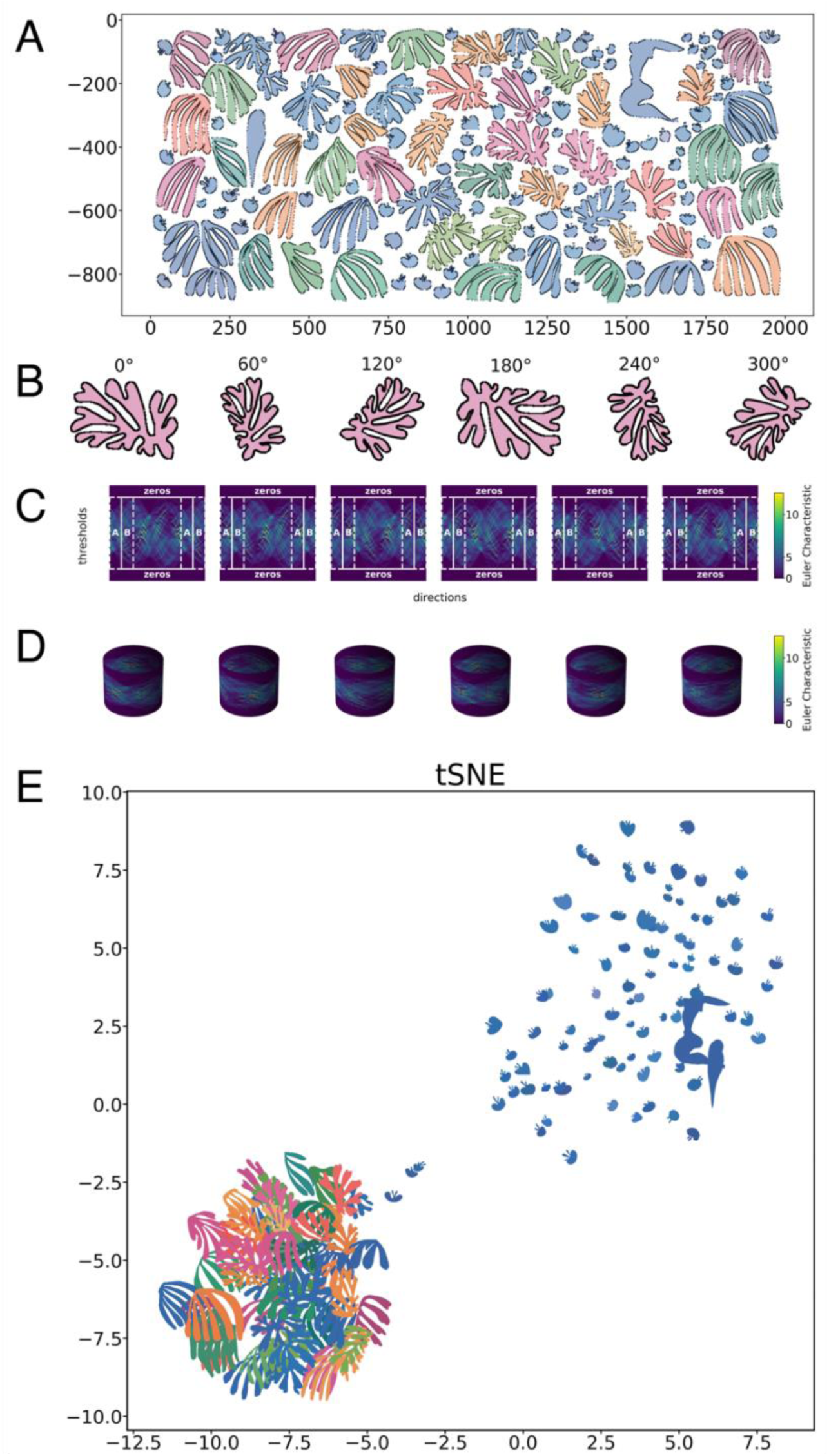
Euler Characteristic Transform (ECT) converts graphs into cylindrical images. A) Graph-representations of cut-outs from *The Parakeet and the Mermaid* (Henri Matisse, 1952). Vertices of each shape graph are plotted in black along the outline. B) Rotations of a shape graph C) The corresponding ECT images for each rotated shape. D) The cylindrical representations of the corresponding ECT images. Cylindrical padding created by corresponding regions "A" and "B" and zero padding are indicated. Euler characteristic (χ) color values are shown. E) Shapes and corresponding colors are projected using t-distributed stochastic neighbor embedding (t-SNE).

**Figure 2:**
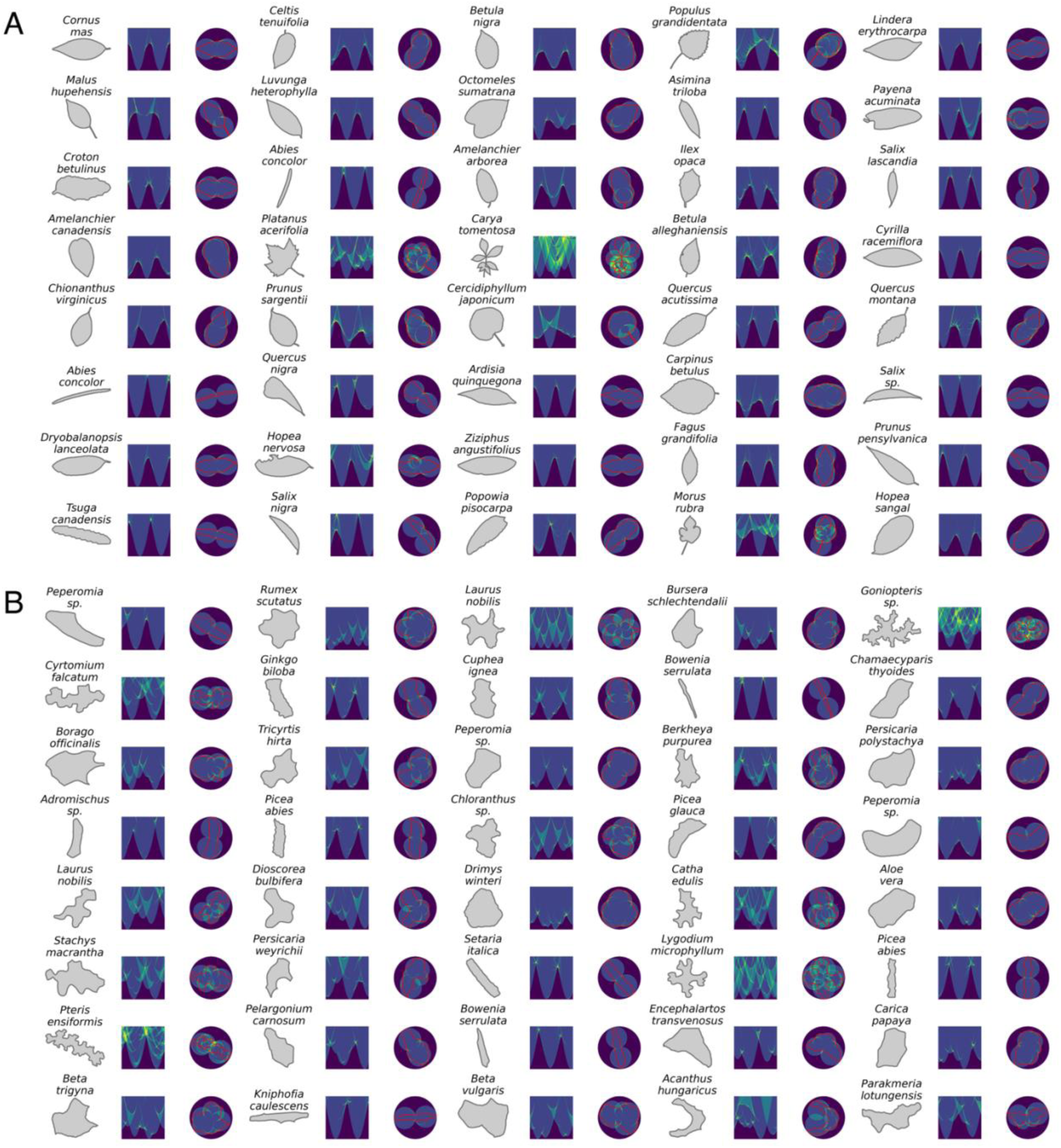
Shapes and associated Euler Characteristic Transform (ECT) images. A) Randomly selected leaf and B) pavement cell shapes and corresponding standard and radial ECT images. Species names and Euler characteristic (χ) color values are provided.

We visualize a wide variety of leaf (**Figure 2A**) and pavement cell (**Figure 2B**) shapes together with their ECT images to better understand the correspondence between these two representations of the same data. These previously published biological shapes represent seed plant leaf shape (Huff et al. 2003; Royer et al. 2005; Peppe et al. 2011; Wang et al. 2024) and vascular plant pavement cell (Vőfély et al. 2019) diversity. All shapes and ECT images are first PCA-aligned and normalized so that they are oriented along their length. The more elongated a shape, the more wave-like its corresponding ECT image appears. This is because, along the bounding circle of the shape, connected components contribute to the Euler characteristic closer to the circumference along the elongated axis and farther along the minor axis. In contrast, more circular shapes produce ECTs with a more uniform band, as connected components contribute evenly across angles and thresholds. Shapes that exhibit greater undulation—through lobing or serration—show elevated Euler characteristic values at thresholds where these features are encountered, often near the center or edge of the shape. By examining (1) the zero-value regions and whether the ECT image appears more wave-or band-like, and (2) the spatial distribution of higher Euler characteristic values, one can infer the shape’s aspect ratio and degree of undulation, which are traditionally measured using circularity or solidity.

### Classification from ECT images

We trained a convolutional classifier to predict shape identity from ECT images for two settings: (i) pavement cell classes and (ii) leaf family or group. The network comprised three to five convolutional blocks (3×3 kernels with batch normalization and nonlinearities), spatial dropout, global average pooling, and a linear classifier with label smoothing; for larger label sets we used class-balanced sampling. Models were optimized with cross-entropy loss and AdamW (Loshchilov and Hutter 2018) using a learning-rate schedule of 1e-3, batch size 512, for up to 50 epochs with early stopping on validation macro-F1 (patience 5). Inputs were normalized per image. We report accuracy, macro-F1, and per-class F1 on held-out test sets and include confusion matrices.

### Approximate inverse segmentation from radial ECT

To assess the extent to which ECT preserves shape information and to provide an interpretable mapping back to geometry, we trained a CNN model to approximate the inverse mapping from radial ECTs to shapes. We first prepared paired inputs and targets from raw leaf outlines provided as 2D coordinate arrays. Coordinates were centered at the origin, oriented consistently using the internal normalization of the ECT framework, and isotropically scaled so that the maximum radial extent equaled 1. The motivation behind using a radial version of the ECT is that now ECT features can be superimposed on top of the original shape mask as an input for subsequent models.

The radial ECT was computed with 360 angular directions and 360 radial thresholds uniformly spanning [0, 1], then rendered as single-channel 256×256 images in polar coordinates. Corresponding binary masks of the normalized shapes were rasterized at 256×256. For qualitative checks, we inspected overlays of ECT images with superimposed outline traces (**Figure 2**).

We employed a U-Net encoder–decoder (Ronneberger et al. 2015) with four down-sampling stages. Each stage used two 3×3 convolutions with batch normalization and ReLU activations (“DoubleConv”), followed by 2×2 max pooling in the encoder, increasing channels from 1 (ECT input) to 64, 128, 256, and 512, with a 1024-channel bottleneck. The decoder used bilinear upsampling with skip connections from the encoder to recover spatial detail and progressively reduced channel depth back to 64. A final 1×1 convolution produced a single-channel logit map.

Training optimized binary cross-entropy with logits optimized using Adam (Kingma and Ba 2017) (learning rate 1×10⁻⁴), batch size 16, for 50 epochs. Data were split deterministically into training/validation/test sets (80/10/10) with a fixed random seed for reproducibility. Augmentations reflected ECT invariances: cyclic horizontal shifts (wrap-around on the angular axis) and mild vertical scale jitter. Predictions were thresholded at 0.5; post-processing retained the largest connected component and filled small interior holes to match ground-truth conventions. Model selection used the best validation Dice coefficient.

### Venation prediction from blade geometry

We constructed a latent morphometric model of grapevine leaves to parameterize joint blade-vein shape variation. Blade and vein landmarks for each leaf were concatenated and subjected to generalized Procrustes analysis to remove translation, scale, and rotation, yielding a mean configuration and Procrustes-aligned shapes. Principal Component Analysis (PCA) on flattened aligned coordinates captured the dominant modes of variation; the number of retained components was bounded by the rank of the data (minimum of sample count and feature dimension). The mean shape, PCA loadings, explained variances, and per-sample scores formed the basis for subsequent synthesis.

To generate training data that span venation diversity while controlling class balance, we synthesized additional leaves in PCA space using a Synthetic Minority Oversampling Technique (SMOTE)-like interpolation within genotype classes. For each leaf we identified k nearest neighbors of the same class and linearly interpolated between the index leaf and a randomly selected neighbor to produce new latent scores until per-class targets were met. Synthetic scores were inverse-transformed to recover separate 2D coordinate sets for blade and vein structures. To promote invariance, coordinates were randomly rotated within [−180°, 180°]. We then computed radial ECTs for blade and vein using a unit bound radius, 360 uniformly spaced directions, and 360 thresholds from 0 to 1. ECT images were rendered as 256×256 single-channel images in polar coordinates and standardized by centering, isotropic scaling, and canonical re-orientation. The affine normalization derived during ECT standardization was applied to the original coordinates so that binary masks rasterized from blade and vein were aligned in the blade’s ECT frame. Visual overlays of outlines on ECTs were used for quality control.

For venation prediction we trained a semantic segmentation model to infer vein masks from blade geometry alone. The network was a U-Net with four encoder–decoder stages operating on two input channels (blade ECT and blade mask) and producing a single-channel vein mask. Each encoder stage comprised two 3×3 convolutions with batch normalization and ReLU activations followed by 2×2 max pooling, increasing channel depth to a bottleneck that preserved high-level context; the decoder used bilinear upsampling with skip connections to recover spatial detail, and a final 1×1 convolution produced logits. Training optimized a composite loss (binary cross-entropy and Dice, equally weighted) with Adam (learning rate 5×10⁻⁴), batch size 16, for 100 epochs, using an 80/20 deterministic split and model selection by best validation Dice.

## Results

### Classification

Although embedded in 2D space, a closed contour outline can be flattened to a 1D vector for representation. For example, chaincode is a lossless compression method that represents contours as 1D vectors (Kuhl and Giardina 1982). Convolutional Neural Networks (CNNs) are particularly suited to image representations. The ECT transforms a closed contour that would otherwise be depicted as a simple shape mask or a 1D lossless compressed vector into a feature rich matrix, or an image. In this image one axis is the directional axis angles and the other thresholds, and the pixel values are the Euler Characteristic values. The ECT encodes topological features into an image, on which CNNs can predict and classify data. Beyond featurizing the closed contour, the ECT presents rotation invariant properties in the context of CNNs. While CNNs can handle translation via the use of a kernel window, they are inherently sensitive to rotation and often learn (and do not intrinsically handle) rotational variance. Because one axis of the ECT matrix represents angles from 0 to 2𝜋, the ECT transforms rotational variance into translational variance. For the classification models described below we tried three different ECT orientations: the ECT calculated on randomly oriented contours, on Principal Component Analysis (PCA)-oriented contours, and on randomly oriented contours with padding that creates a cylinder representation of the ECT in the eyes of a CNN (**Figure 1B-D**; McGuire 2024). These different methods of preparing the ECT for CNNs performed equivalently (data not shown) on our data, and we only present results for PCA-oriented contours. However, this does not mean that the rotational invariant properties conferred by ECT are not useful for other data types.

We first built a CNN-ECT classifier on 172,790 leaf shapes (Li et al. 2017; Wang et al. 2024) representing 14 classes sampled in depth from various crops and plant species and 2 classes representing diverse plant species with distinct leaf shapes across seed plants (“Leafsnap” and “Transect” classes). We observe extremely high prediction performance with leaf classification from ECT images achieving 84.6% accuracy and macro-F1 of 0.797 on a held-out test set (n = 34,271 across 16 classes, per class results shown in **Table 1**, **Figure 3**). Macro average precision was 0.899.

**Figure 3:**
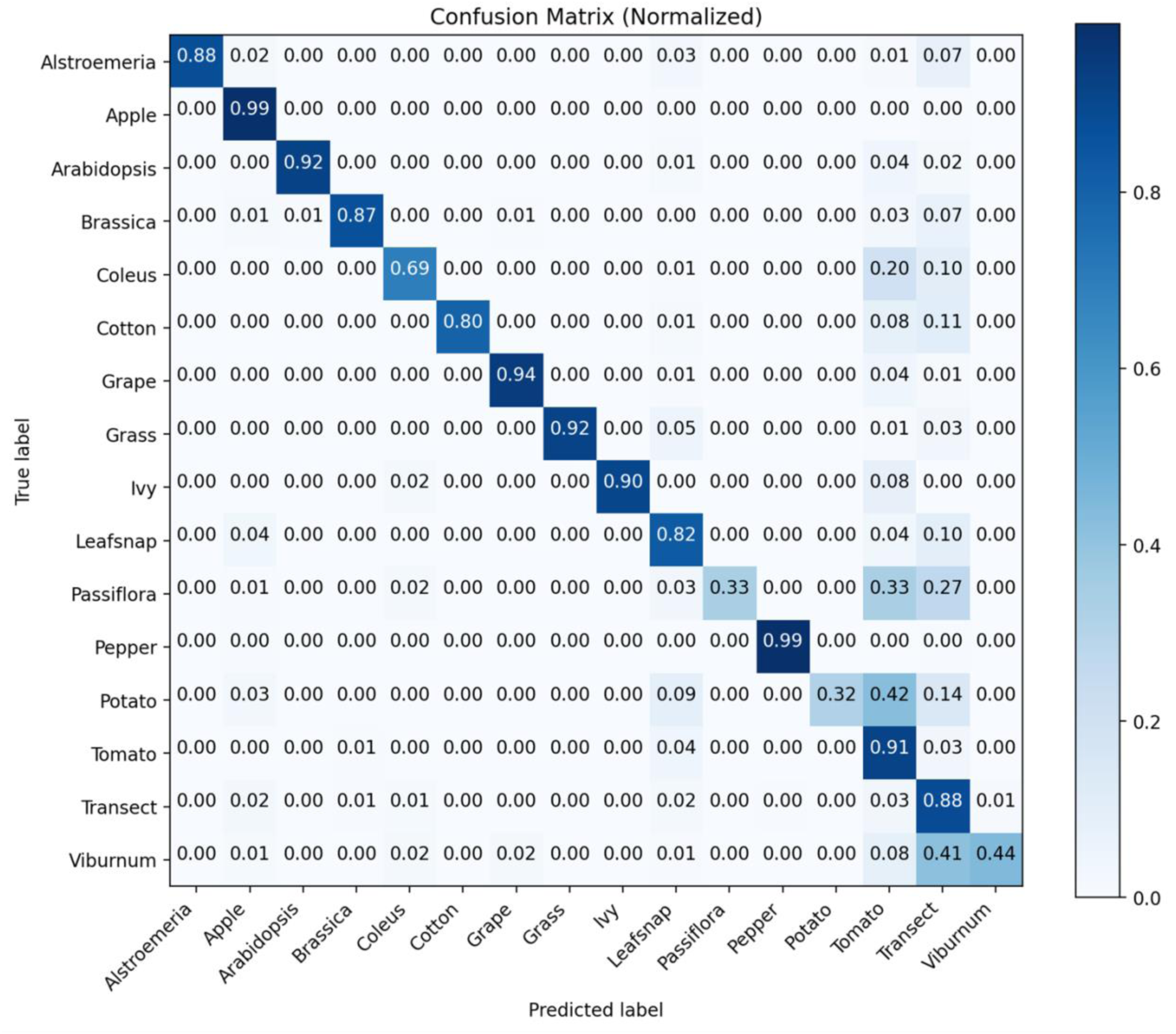
A confusion matrix showing model predictions vs. true labels for a CNN classifying leaf shapes from 14 plant species sampled in depth and two groups (Leafsnap and Transect) representing diverse leaf shapes across seed plants. Rows represent true class labels and columns represent predicted labels; cell values indicate the number of samples in each category. Cells along the main diagonal represent correct classifications.

**Table 1:**
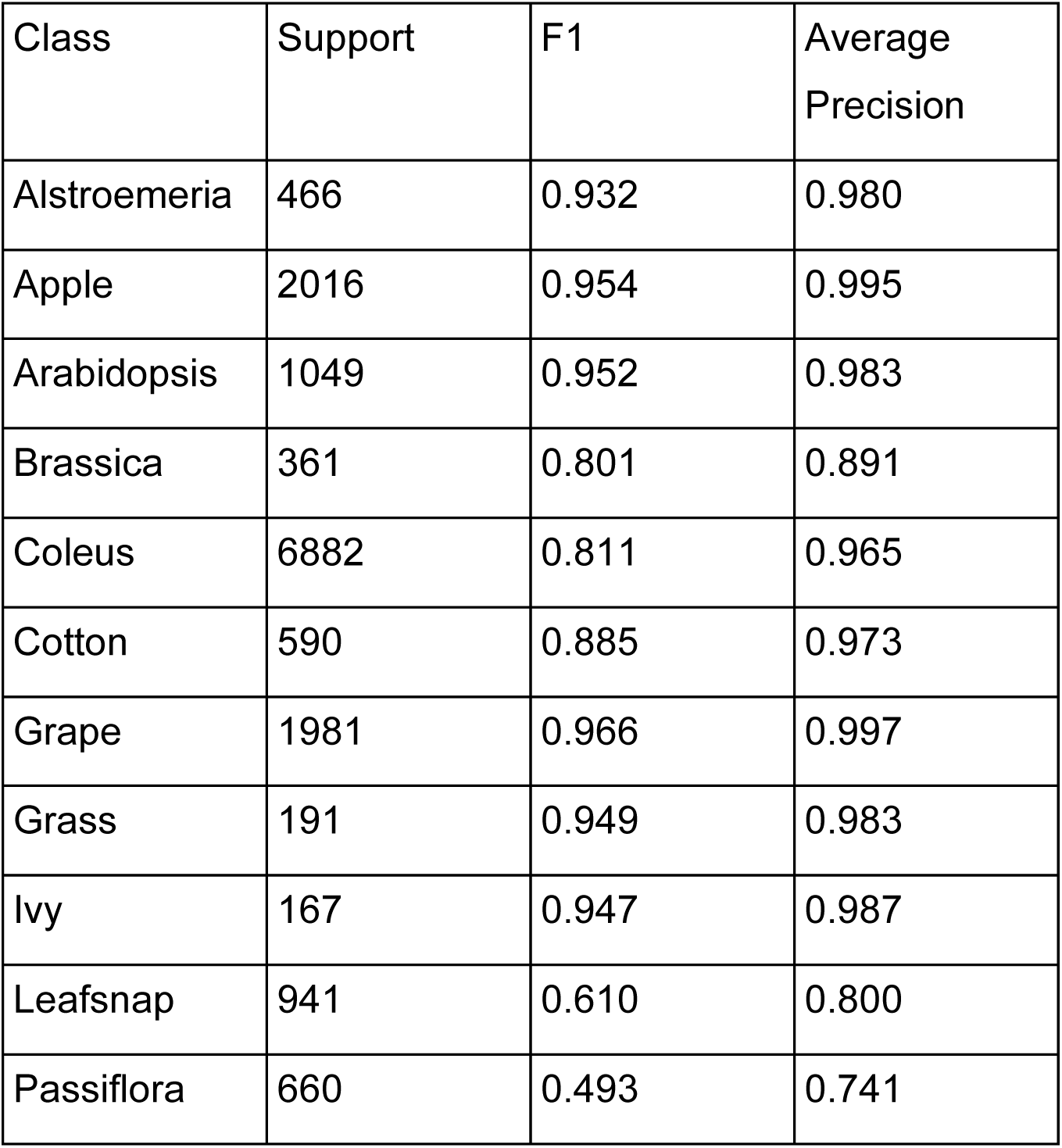
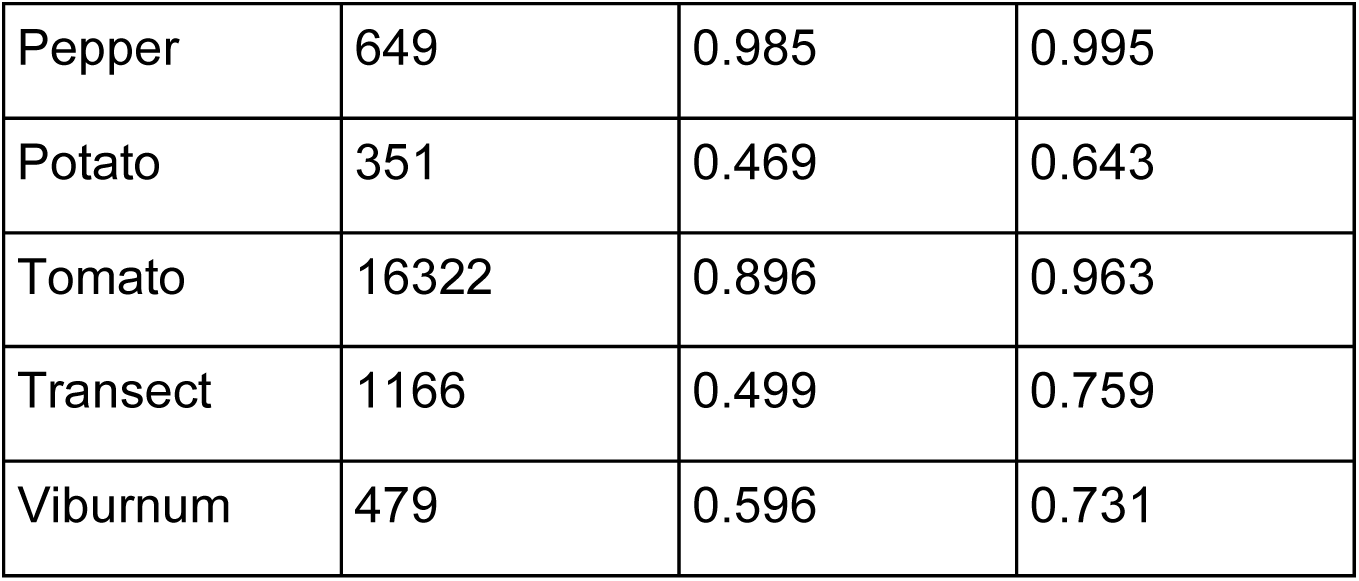
Classification results for sampling of 14 plant species and 2 diverse classes (Leafsnap and Transect)

We next analyzed the more complex and irregular shapes lacking orientation of 11347 pavement cells from four classes (ferns, gymnosperms, monocots, and eudicots; Vőfély et al., 2019). The performance is less than that observed for leaves of disparate shape classes, reflecting the complexity and irregularity of these shapes, yet still the model shows successful classification with 67.4% accuracy and macro-F1 of 0.612 on a held-out test set (n = 998 across four classes, **Table 2**, **Figure 4**). Given the relative performance across leaf shape families of 51.2% (see below), we interpret this level of accuracy, on broad plant families using complex shapes, to be very good. Performance tracked class imbalance, with highest F1 for ferns, moderate for monocots and eudicots, and lowest for gymnosperms. Macro average precision was 0.622.

**Figure 4:**
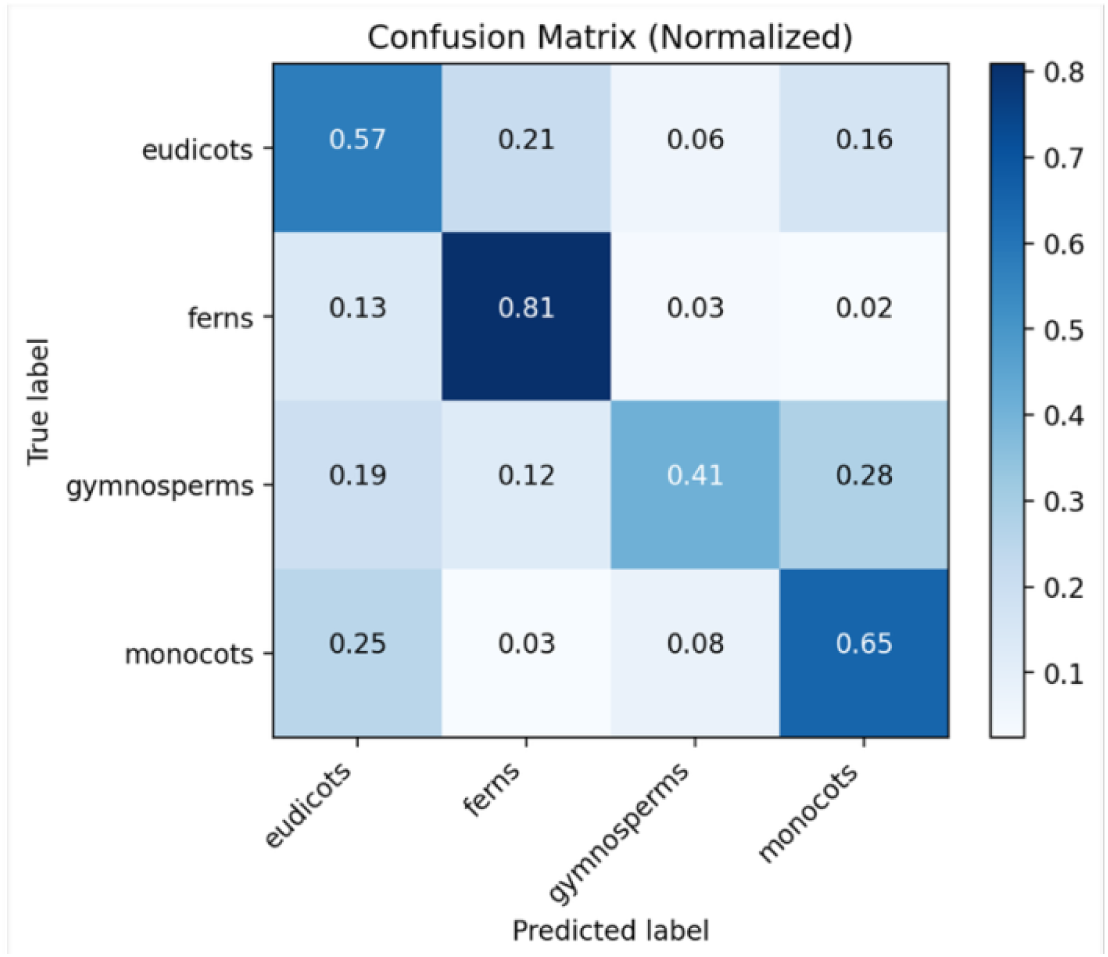
A confusion matrix showing model predictions vs. true labels for a CNN classifying pavement cell shapes into four plant taxonomic groups (eudicots, ferns, gymnosperms, monocots) in a test set of 998 samples. Rows represent true class labels and columns represent predicted labels; cell values indicate the number of samples in each category. Cells along the main diagonal represent correct classifications.

**Table 2:** Classification results for 4 classes of pavement cells.

| Class | Support | F1 | Average Precision |
| --- | --- | --- | --- |
| eudicots | 249 | 0.550 | 0.532 |
| ferns | 449 | 0.822 | 0.892 |
| gymnosperms | 115 | 0.459 | 0.445 |
| monocots | 185 | 0.619 | 0.621 |

To test the performance of ECT in aiding CNN classification, we next turned to a much larger and highly diverse dataset. Using LeafMachine2 to automatically extract leaf shapes from herbarium vouchers (Weaver and Smith 2023), we classified on over 4 million leaves across 11 plant families. Previously classifying on over 10,000 leaves across hundreds of plant families, we achieved classification rates of 10.2% using simple shape descriptors alone, 27.3% using a persistent homology approach, and 29.1% combining the two methods (Li et al., 2018). On this much larger and more challenging dataset, the ECT-based model still achieves meaningful classification. LeafMachine2 herbarium leaf classification (11 families) on a held-out test set achieved 51.2% accuracy and macro-F1 of 0.356 (n = 813,619, **Table 3**, **Figure 5**). Macro average precision was 0.382.

**Figure 5:**
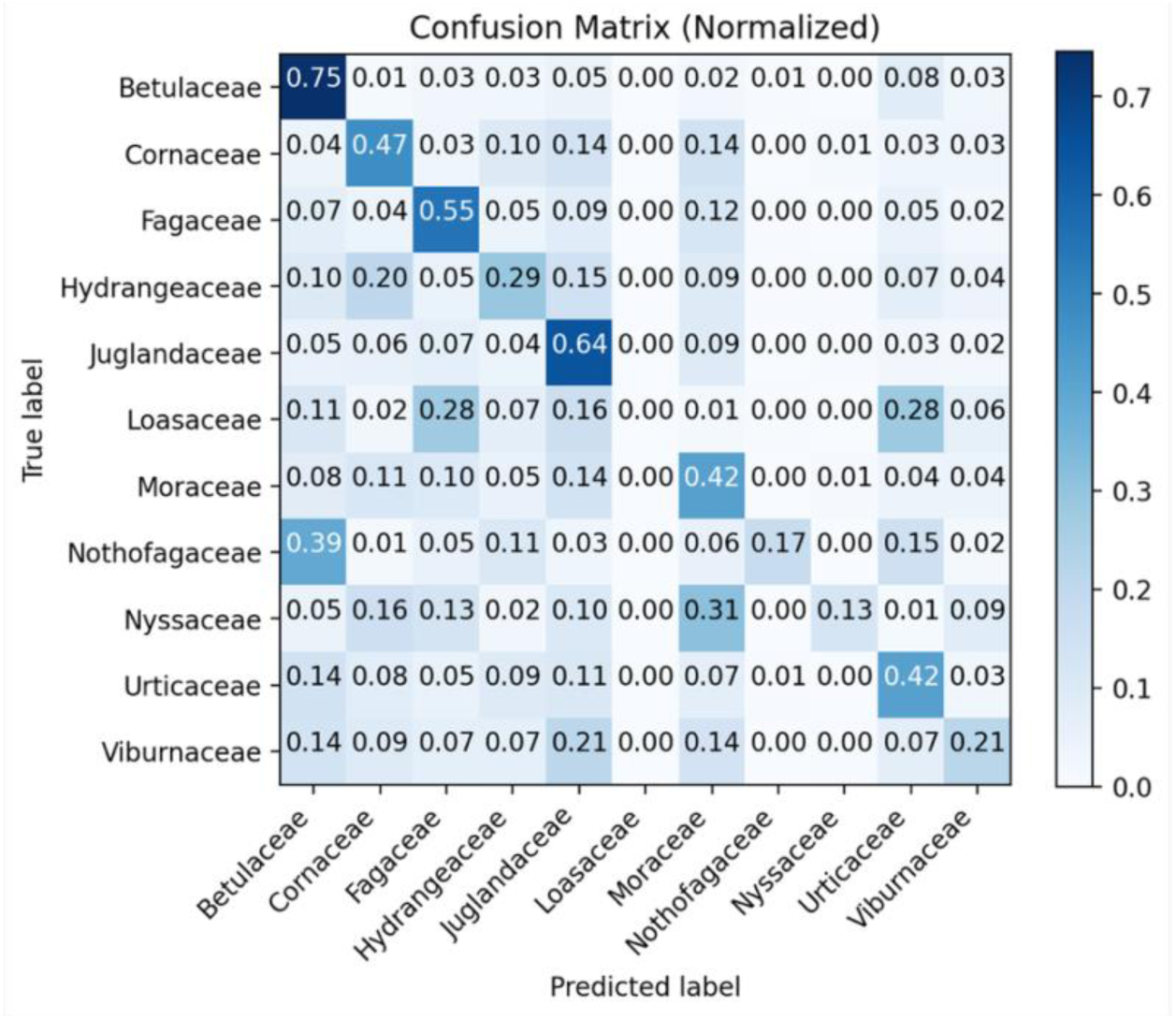
A confusion matrix showing model predictions vs. true labels for a CNN classifying over 4 million leaf shapes from 11 plant families. Rows represent true class labels and columns represent predicted labels; cell values indicate the number of samples in each category. Cells along the main diagonal represent correct classifications.

**Table 3:** Classification results on over four million leaves from 11 classes.

| Class | Support | F1 | Average Precision |
| --- | --- | --- | --- |
| Betulaceae | 193582 | 0.732 | 0.834 |
| Cornaceae | 58323 | 0.420 | 0.417 |
| Fagaceae | 179013 | 0.634 | 0.754 |
| Hydrangeaceae | 37313 | 0.246 | 0.201 |
| Juglandaceae | 97992 | 0.523 | 0.548 |
| Loasaceae | 610 | 0.000 | 0.040 |
| Moraceae | 57784 | 0.316 | 0.295 |
| Nothofagaceae | 6701 | 0.226 | 0.189 |
| Nyssaceae | 9875 | 0.177 | 0.149 |
| Urticaceae | 39740 | 0.327 | 0.312 |
| Viburnaceae | 132686 | 0.314 | 0.460 |

### The radial ECT and approximate inverse using semantic segmentation

For classification, we describe and use the ECT as a 2D matrix, defined by axes of angles and thresholds. The Euler characteristic values in this matrix are topological features that are defined by features in the original contour. Additionally, this matrix is approximately centrosymmetric, reflecting the directional sampling of thresholds across the sampled axes. To create a spatial correspondence between the features of the ECT and the original shape mask from which it is derived, we construct the radial ECT by converting the top half of the ECT matrix into polar coordinates (See **Figure 2**). This radial ECT representation can then be projected onto the original contour, demonstrating how ECT features and original contour features correspond.

A semantic segmentation CNN assigns class identity to each pixel in an image. The proof of Turner et al. (2014) demonstrates that there is a one-to-one function between data objects and their ECTs which mathematically implies that an inverse function for the ECT exists, with some available options in practice (Fasy et al. 2025). However, efficient computation of such an inverse function remains elusive. Because features in the radial ECT correspond to the original contour, we hypothesized that we would be able to train a semantic segmentation CNN to predict the original shape mask of a contour using its radial ECT as an input. Using over 10,000 leaves collected from across the flowering plants (Li et al. 2017; Wang et al. 2024), we trained a U-Net model to predict the original shape mask from its radial ECT image. Visually, the predicted masks correspond closely to the original leaf shapes (**Figure 6**). The held out Dice scores similarly reflect high prediction performance of 97.5% after 50 epochs of training. Although not a true inverse function, we demonstrate that semantic segmentation using a CNN can approximate the inverse of the ECT.

**Figure 6:**
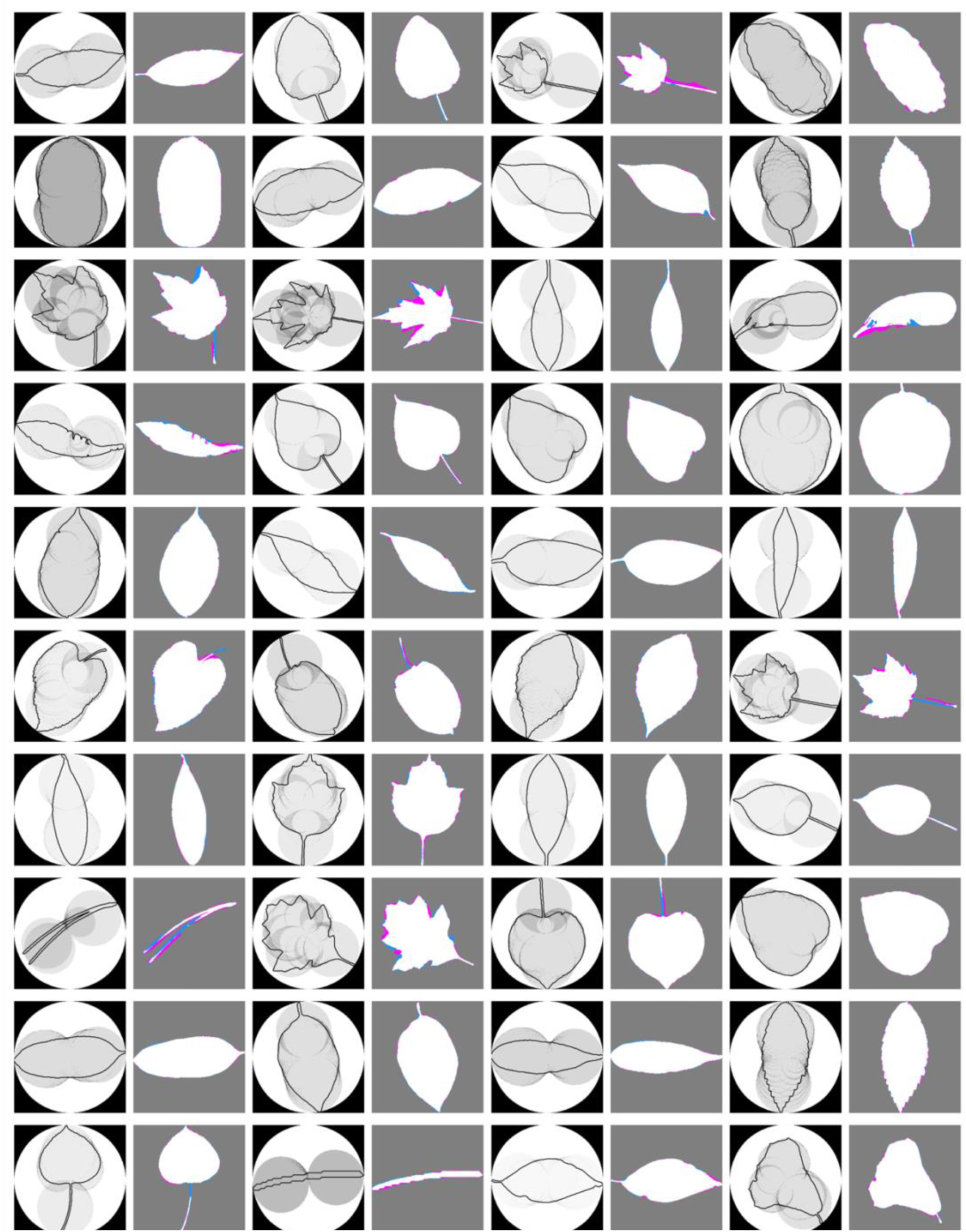
Prediction of segmented leaf masks from corresponding radial ECTs using a CNN. For 40 randomly selected leaves, their radial ECT with superimposed leaf outline (left) and CNN predicted pixels of the leaf shape mask (right), with true positives (white), false positives (magenta), false negatives (dodgerblue), and background (gray) indicated by color.

### The leaf blade contains sufficient information to predict the vasculature

The success of predicting the original shape mask from a spatially corresponding radial ECT has important implications. From one set of embedded features in an image, is it possible to predict another, given a mapping between the two? Such a scenario is present in leaves themselves. The contour of the blade that we have been examining thus far is embedded with the vasculature in the same leaf. We hypothesized that by using semantic segmentation, we would be able to predict the vasculature of the leaf from information in the blade. We turned to grapevine leaves and the field of ampelography to answer this question, due to the fact that every grapevine leaf has five major veins, permitting Procrustean-based geometric methods. We used high resolution landmark data of leaves with outlines of the blade and vasculature (Chitwood et al. 2024). We used a Synthetic Minority Oversampling Technique (SMOTE; Chawla et al. 2002) approach to augment data within classes and create a sufficient number of synthetic leaves to train a model. Using the shape mask of the blade and its corresponding radial ECT, a semantic segmentation U-Net model was trained to predict vein pixels using only information from the blade (**Figure 7**). The fine, intricate structure of the vasculature is reasonably predicted. Because veins are thin, elongated structures, a portion of false positives reflect “near misses” that still reconstruct a complete and connected vascular system. This is reflected in the performance statistics with the U-Net achieving a peak validation Dice coefficient of 0.583 over 100 training epochs, with performance plateauing near 0.57–0.58 after approximately epoch 70. The moderate Dice score is consistent with the inherent difficulty of thin-structure segmentation, where single-pixel offsets along vein boundaries disproportionately penalize the metric relative to the biological fidelity of the reconstruction. We conclude that just as the mapping between a contour and its radial ECT allows for the approximation of the inverse, that similarly a unique mapping between tissues in the leaf allows vasculature to be predicted from the blade.

**Figure 7:**
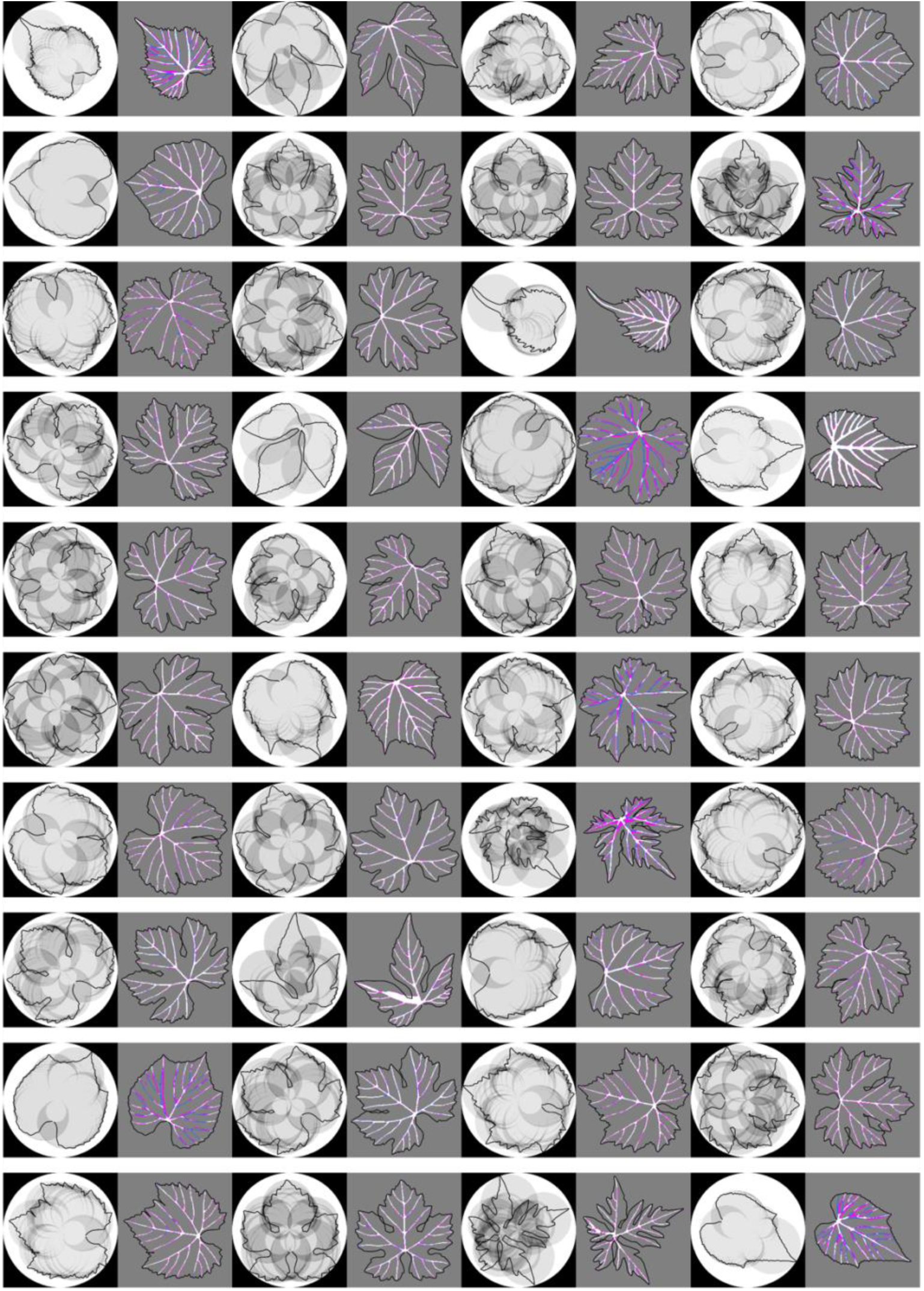
Forty randomly selected synthetic leaves are shown in paired panels. Left panels display the blade Euler Characteristic Transform (ECT) input in inverted grayscale with the leaf blade outline overlaid. Right panels show pixel-level agreement between predicted and ground-truth vein masks: true positives (white), false positives (magenta), and false negatives (blue); gray regions indicate true negative background pixels.

## Discussion

Under the conditions described by Turner et al. (2014), the ECT gives a unique signature for embedded graphs, meaning that different graphs produce different transforms. By transforming data objects into graphs (from closed contours representing leaf outlines, in this case) we permit the use of this tool that allows us to readily sample all available information contained within. The matrix representation of the ECT lends itself to vector-based representations, and as we have discussed, treating it as a literal image. This opens the door for representing topological structure of a shape as an image that can immediately be used within a neural network framework. As we showed with leaf shapes representing distinct shape classes (**Table 1**), the irregular shapes of pavement cells (**Table 2**), and with a very large number of leaves representing shape diversity across plant families (**Table 3**), not only does the ECT provide a convenient image based summary of data, but it also featurizes the data within a topological framework that complements the original data structure. We have also investigated the use of the “radial ECT”, where the top half of the ECT is visualized in a polar image which makes obvious the geometric connection to the original input shape. We note that because of the simplicity of the inputs (each leaf outline is a closed contour, which is topologically equivalent to a circle), duality results from topology imply that the entirety of the ECT is encoded in this radial ECT even though it only shows half of the original input image.

The mathematical properties of the ECT make this construction an exciting addition to the plant morphology toolkit. The uniqueness of the representation (when using enough directions) lends itself to being an excellent summary of the input data, while still being in a vector form that is easy to pass to any deep learning or machine learning frameworks. Because the representation is unique, in theory one should be able to reconstruct an input structure from the ECT. While research is ongoing to do this in exact cases, we feel that using the CNN as done here opens the door to an easier to access approximation of the inverse map that will be very useful in practice. Further, the ECT has an inherent connection to the rotation of the inputs: specifically, rotation of an input shape corresponds to translation of the output ECT. Future iterations of this construction and deep learning pipelines could be used for additional alignment and symmetry tasks by harnessing this association.

We have also used this approximation of the inverse map to test connections with biological reality as well. Within leaves there are two closed contours: one representing the blade and the other the intricate vascular system embedded within it. Just as the mapping allows us to predict the original data from its ECT, so too the biological mapping (reflecting developmental constraint) between blade and vein allows us to predict the vascular network of grapevine leaves using only information from the blade alone (the original shape mask and the corresponding ECT). This creates a unique, mathematical reinterpretation of the classic phenomenon of developmental constraint, that there is a mapping across tissues within the leaf and that all the information to predict the morphology of one embedded tissue type can be found in the other. That ultimately the totality of phenotype (all traits) arises pleitropically from a single genotype (the collection of allelic variants in a genome) reflects how constraint operates and that there would be unique mappings between the different parts that comprise an organism.

The ECT poses both opportunities and challenges with scalability of shape analysis. As an example to demonstrate the usefulness of the ECT in this paper, we analyzed over 4 million leaf shapes from 11 plant families extracted from herbarium vouchers with LeafMachine2 (Weaver and Smith 2023) encoded as embedded graphs. However, this is only a small sampling (orders of magnitude smaller than the ultimate number) of the leaf shapes that will ultimately be extracted. Future work related to improved storage and access of this particular class of inputs could yield further insights by allowing biological researchers to interface with even larger datasets. In addition, mathematically there is no restriction on the dimension of the input shapes or the dimension in which the data is embedded. For the input shape dimension, graphs can be considered as 1-dimensional input objects since they are made up of lines; future versions of this work including information from higher dimensional constructions such as simplicial complexes hold the potential to find even more structure depending on the input data.

For the dimension in which the data is embedded, here we have only looked at 2D image data. In previous work we have evaluated 3D scans of barley shapes using a simplified version of the machine learning pipeline (Amézquita et al. 2022); however, extending this work to 3D (for example on 3D scans, protein structures, canopy and root architectures, or other higher dimensional structures) could yield additional structural information if the curse of dimensionality can be overcome; see e.g. (Paik 2023).

The ECT thus establishes itself as a powerful tool that efficiently maps complex, high-dimensional biological data into a deep-learning-compatible image format while preserving an essential property: topological injectivity. This injectivity, demonstrated through the successful approximation of the inverse function and the prediction of complex venation from simple blade geometry, confirms that the ECT is not merely a feature descriptor but a faithful, compressed representation of the data’s topological and geometric information. By overcoming the twin challenges of shape variance and deep learning’s rotational sensitivity, the ECT provides a rich and robust representation for analyzing and storing any graph-representable biological structure, from 2D cellular networks to the countless higher-dimensional protein and macromolecular assemblies that now define modern structural biology.

Our results offer a novel, mathematically grounded interpretation of developmental constraint, demonstrating that the information necessary to construct a complex tissue (the vasculature) is predictably encoded within the geometry of another (the blade outline). This success, alongside the unprecedented classification accuracy on massive, diverse datasets, shows that the ECT is an essential tool for modern plant morphology with the potential to transition from descriptive morphometrics to a truly predictive and integrated analysis of development, evolution, and constraint across all scales of biological complexity.

## Acknowledgements

YA, AYH, RV, EM, and DHC are supported by the National Science Foundation Plant Genome Research Program award numbers IOS-2310355, IOS-2310356, and IOS-2310357. RV and DHC are supported by MSU AgBioResearch.

## Author contributions

YA, SM-S, SP, WNW, NK, and WY developed essential methods, machine learning models, or data to apply the Euler Characteristic Transform (ECT). KA-P, AB, JCC, JCL, BAD, DVE-M, QF, KMG-L, JNG-C, SG-R, CNG, GEG-L, JCH, CLH, ATH, JJH-O, SK, JLennon, ZL, JLi, BL, JLin, PL-S, ML-A, CM-M, AM-L, NAM, IAO, OP-F, CP-R, EP-H, CP, LS, SS, FT, JAT, MAT, JKT, JJVW, DV-H, EXW, NW, and JZ provided conceptualization and exploratory data analysis to apply the ECT to plant data. BLM, AR-H, EBJ, ZM, LD-G, AYH, SAS, ZL, RV, AR-C, EM, and DHC provided funding, instruction, supervision, and guidance. YA, EM, and DHC developed figures, wrote, and finalized the manuscript. All authors read, edited, and approved the manuscript.

## Data Availability Statement

Data and code to fully reproduce the results of this manuscript are available in the following repositories:

Training and Evaluation code: https://github.com/yemeen/ectnet

Leaf shape outlines as .npy files from Li et al., 2017: https://figshare.com/articles/dataset/Modified_leaf_shape_contour_data/25435936

LeafMachine2 outlines: https://zenodo.org/communities/leafmachine2/records?q=&l=list&p=1&s=10&sort=newest

Pavement cell outline as .npy files from Vőfély et al., 2019: https://datadryad.org/stash/dataset/doi:10.5061/dryad.g4q6pv3

